# Sequence and primer independent stochastic heterogeneity in PCR amplification efficiency revealed by single molecule barcoding

**DOI:** 10.1101/011411

**Authors:** Katharine Best, Theres Oakes, James M. Heather, John Shawe-Taylor, Benny Chain

**Affiliations:** Division of Infection and Immunity, UCL, London; CoMPLEX, UCL, London; Department of Computer Sciences, UCL, London

## Abstract

The polymerase chain reaction (PCR) is one of the most widely used techniques in molecular biology. In combination with High Throughput Sequencing (HTS), PCR is widely used to quantify transcript abundance for RNA-seq and especially in the context of analysis of T cell and B cell receptor repertoires. In this study, we combine molecular DNA barcoding with HTS to quantify PCR output from individual target molecules. Our results demonstrate that the PCR process exhibits very significant unexpected heterogeneity, which is independent of the sequence of the primers or target, and independent of bulk experimental conditions. The mechanistic origin of this heterogeneity is not clear, but simulations suggest that it must derive from inherited differences between different DNA molecules within the reaction. The results illustrate that single molecule barcoding is important in order to derive reproducible quantitative results from any protocol which combines PCR with HTS.

## Introduction

The efficiency of a PCR reaction is known to vary widely, depending on many different factors. These include the properties of the primers (1–3), the sequence to be amplified (4), in particular the GC content (5, 6) as well as the reaction conditions and type of polymerase. If one wishes to quantify the amount of a given template by PCR (qPCR) the general approach is to compare an unknown sample to a dilution series of standards, on the assumption that all variables remain the same between sample and standard and hence PCR efficiency remains constant.

The introduction of high throughput sequencing (HTS) (7, 8), in which many DNA molecules are sequenced individually in parallel, allows the possibility of quantifying many DNA molecules simultaneously by counting the number of times the sequence for each molecule occurs in a sequence run. This approach forms the basis for RNA-seq, in which transcript abundance is measured by sequencing cDNA libraries, and counting the number of sequences mapping to each transcript. An extension of this approach is the analysis of the antigen-specific receptor repertoire by sequencing cDNA or genomic samples of B or T lymphocytes, and counting the number of times each different receptor is identified (9). Most current parallel sequencing technologies require nanomolar amounts of starting material (typically > 10^10^ molecules), even when the output of the reaction may only be in the order of 10^7^ molecules (for example the Illumina MiSeq). In order to achieve this amount of starting material some degree of PCR amplification is usually required. This is especially true when the amount of starting material may be extremely small, for example in the case of single cell RNA-seq (10). The reproducibility of the PCR amplification process therefore becomes a key factor.

The use of molecular barcodes provides one approach to dealing with single molecule quantification and mitigating the effects of PCR heterogeneity. A library of diverse short DNA sequences (barcodes or tags) are introduced into the sequences to be analysed at an early step in the protocol, in such a way that each target molecule incorporates a different tag which remains associated with it throughout the amplification protocol. The barcodes can be introduced during a reverse transcription step, or by ligation. For instance, Miner et al (11) and McCloskey et al (12) both ligate nucleotide sequences to uniquely label initial DNA target molecules to identify sequencing redundancy as well as using batch stamps to identify sequencing contamination from other samples. In the work of Casbon et al (13) degenerate base regions are ligated to each DNA fragment to assess whether observed differences between sequence reads are true variants or sequence error. Kivioja et al (14) apply a similar unique molecular identifier technique to human karyotyping. Mamedov et al (15) and Shugay et al (16) use barcodes to provide PCR and sequencing error correction of TCR repertoires.

In this study we use a protocol that incorporates ligation of a random 12 base pair-long nucleotide barcode to uniquely label each individual molecule. We use this protocol to investigate the extent of variation in PCR amplification. In order to rigorously assess the possible sources of this heterogeneity we develop an efficient PCR simulator, which incorporates both amplification and sampling heterogeneity, with which to compare our experimental results. The PCR amplification is an example of a branching process, and although there has been considerable theoretical work on such processes, the complexity of heterogeneous branching processes makes analytical modelling challenging in most realistic examples (17–19). Detailed models of the physical parameters involved in the PCR cycles have been developed, to answer questions about the probability of replication in an individual cycle or the evolution of the population over a number of cycles (20–24). Additionally mathematical models have been used to investigate the error profile in PCR protocols (25) or the presence of non-targeted product through nonspecific priming (26). The increase in computing power has made it feasible to develop PCR simulations using realistic numbers of starting molecules, with reasonable run times. The model we describe includes both an amplification step and a sampling step to simulate the typical workflow of an RNA-seq experiment.

Our results demonstrate that the PCR process exhibits very significant unexpected heterogeneity, which is independent of primers, target sequence or bulk experimental conditions. The mechanistic origin of this heterogeneity is not clear, but simulations suggest that it must derive from inherited differences between different DNA molecules within the reaction. The results illustrate that single molecule barcoding is essential in order to derive reproducible quantitative results from any protocol which combines PCR with HTS.

## Methods

### Ethics

This study was approved by the joint UCL/University College London Hospitals NHS Trust Human Research Ethics Committee and was carried out in accordance with relevant guidelines and regulations. Written informed consent was obtained from all participants (University College Hospital 06/Q0502/92)

### Sample collection

For TCR repertoires sequenced from whole peripheral blood (PB), 2.5ml of healthy adult volunteer PB was drawn into Tempus tubes and RNA was extracted following the manufacturer’s instructions (Life Technologies). Residual DNA was removed using the TURBO DNase kit, and globin mRNA was depleted using GLOBINclear (both Life Technologies).

Alternatively, for TCR repertoires sequenced from purified peripheral blood mononuclear cells (PBMC), 60ml of healthy adult volunteer peripheral blood was drawn and PBMC were isolated through density-gradient centrifugation using Ficoll-Paque PLUS (GE Healthcare Lifescience). RNA was isolated using an RNeasy Mini Kit (Qiagen).

For monoclonal sequencing experiments, RNA was isolated (RNeasy Mini Kit, Qiagen) from a T cell clone (KT2) specific for tetanus toxoid antigen (30). 1μg of RNA was treated with RQ1 DNase (Promega) following manufacturer’s instructions to remove any residual genomic DNA.

### TCR amplification and sequencing

RNA was reverse transcribed using oligos directed against the 5’ region of TRAC and TRBC (αRC2 and βRC2). All primers are from Sigma-Aldrich and sequences can be found in Supplementary information. 11μl of DNase treated RNA were mixed with 0.5μM αRC2, 0.5μM βRC2 and 0.5mM of each dNTP (Invitrogen) to total 19.5μl, and then incubated at 65°c for 5 min and cooled rapidly on ice for > 1 min. 1× FS buffer (Invitrogen), 5mM DTT (Invitrogen), 30-60 units RNasin Ribonuclease Inhibitor (Promega) and 300 units SuperScript III reverse transcriptase (Life Technologies) were added before incubation at 55°c for 30 min in a total volume of 30μl. 40mM NaOH were added to remove any remaining RNA and the sample was incubated at 70°c for 15 min. 0.5M sodium acetate were added to adjust the pH before the reverse transcription product was purified using MinElute columns (Qiagen).

cDNA underwent a 5’-RACE ligation labelling step, to ligate an oligo containing a random nucleotide barcode and Illumina sequencing primer 2 (6N_I8.1_6N_I8.1_SP2), for PBMC and KT2 or containing the first half of the random nucleotide barcode and the sequencing primer for PB samples (T4DNA_6N_SP2). 5μl of cDNA were mixed with 50% PEG 8000 (New England Biolabs), 1× T4 RNA ligase buffer (NEB), BSA (Promega), 1mM hexamine cobalt chloride, 0.33mM ATP (NEB), 0.33μM ligation oligo and 20 units T4 RNA ligase 1 (NEB). Mixes were incubated at 16° for 23 hours followed by a 10 minute heat inactivation step at 65°. Samples were purified using AMPure XP SPRI beads (Appleton Woods) 1:1 following manufacturer’s instructions and eluted in 30-35μl water.

TCR from the PBMC and KT2 samples were amplified in two consecutive PCR reactions. In PCR1 the TCRs were amplified from the constant region of the TCR α- or β-chains (αRC1, βRC1.1 or βRC1.2) to the Illumina sequencing primer SP2 which was added via the ligation. 31μl of ligation product were added to 1× HF buffer (NEB), 0.5μM of αRC1, βRC1.1, βRC1.2 and SP2, 0.5mM of each dNTP (Life Technologies) and 1 unit Phusion (NEB). The 50μl reactions were run on a Thermal cycler: initial cycle 98°C for 3 min; cycle 1-13: 98°C for 15 sec, 69°C for 30 sec and 72°C for 40 sec; final cycle: 72°C for 5 min. The PCR product was purified with AMPure beads and eluted in 30μl water.

In PCR2 the sample was split and the α- and β-chains were amplified separately. During this reaction two index sequences are introduced either side of the molecule (I-X on the constant region end or LX on the V/SP2-region end) in order to multiplex different samples in the sequencing reaction. A second Illumina primer (SP1) and random hexamers were also added on to the constant region end. The sequencing adapters P5 and P7 allow the construct to bind to the Illumina flow cell and were introduced either end of the amplified TCRs. A qPCR reaction was performed (ABI, Applied Biosciences) suing the sample heat cycles as in PCR1. The 25μl reaction contained 13μl PCR1 product, 1× HF buffer (NEB), Cybrgreen, 0.25μM of each dNTP, Rox, 0.05μM SP1-6N-I-X-αRC1 or SP1-6N-I-X-βRC1.1+1.2, 0.5μM SP1-SP5, 0.5μM P7-LX and 0.5 units Phusion (NEB). The qPCR was run until the products reached the Ct value (about 8 cycles). PCR2 product was purified with AMPure beads and eluted in 30μl water.

As mentioned before for the PB samples six random nucleotides were added during the ligation. The other six random nucleotides were added to the constant region end in a single round PCR in order to label every molecule uniquely. For that, the complementary strand (second strand) to the ligated cDNA had to be made first. The AMPure bead purified ligation product was incubated with 1× HF buffer, 0.5μM SP2 primer, 0.5mM of dNTPs and 1 unit Phusion in a 50μl reaction at 98°C for 3 min, lowered slowly (1°C/sec) to 80°C, held at 80°C for 10 sec, lowered slowly (1°C/sec) to 58°C and held at 58°C for 30 sec. After the final extension at 72°C for 1 min the second strand was purified with AMPure beads and a third strand was synthesised using the same conditions as for the second strand. To introduce a random hexamer the following primers were used instead of SP2: SP1-6N-I-X-αRC1 for the α-chain and SP1-6N-I-X-βRC1.1/1.2 for the β-chain.

PB samples are amplified in two different consecutive PCR reactions. In the first PCR the P5 and P7 adapters were introduced (1× HF buffer, 0.5μM P5-SP1, 0.5μM P7-LX, 0.5mM dNTPs and 1 unit Phusion) at: initial cycle: 98°C for 3 min, slowly ramped to 69°C for 15 sec and 1 min at 72°; cycle 2-4: 98°C for 10 sec and 72°C for 1 min; final cycle: 72°C for 5 min. After bead purification the samples were amplified in a second PCR (1× HF buffer, 0.5μM P5s (details below), 0.5μM P7 (details below), 0.5mM dNTPs and 1 unity Phusion) at: initial cycle 98°C for 3 min; cycle 1-24: 98°C for 10 sec, 69°C for 15 sec, 72°C for 40 sec; final cycle: 72°C for 5 min. PCR1 and PCR2 were performed in 50μl reactions. PCR2 products were bead purified and eluted in 30μl water.

Final amplicon products from all sample types were quantified on a Qubit fluorometer (Life Technologies) and sized on a Bioanalyzer (Agilent). Up to 12 samples (at a concentration of 4nM) were multiplexed and sequence on an Illumina MiSeq, using version 2 chemistry 2×250PE kits.

### Data analysis

The FASTQ files produced on the MiSeq were demultiplexed based on the indexes added through PCR and analysed using a modified version of Decombinator [REF]. Decombinator categorises each TCR sequence read by identifying its constituent V gene and J gene, along with the number of nucleotide deletions from each and the non-germline junctional nucleotides inserted during TCR recombination. The five-part Decombinator classifier (DCR) is given by (V gene used, J gene used, number of V deletions, number of J deletions, inserted nucleotides). The modified version of Decombinator used in this study outputs the DCR along with information about the random nucleotide barcode and sequence quality in each sequence read.

The Decombinator output is then passed into a PCR: and sequencing-error correction script. This script first filters sequence reads to remove those where the barcode or sequence quality are poor. It then collects all sequence reads according to their barcode, grouping together those DCRs that appear with identical barcodes. If more than one distinct DCR appears with the same barcode, we take the DCR with the most copies to be the true sequence with that barcode, and the others are aggregated into the largest DCR if they are clearly the product of sequencing error or discarded otherwise. Next, the set of different barcodes associated with the same DCR is considered. Barcodes that are similar and are observed in the context of the same DCR are considered to be derived from the same initial molecule and are therefore aggregated. The size of the set of distinct barcodes found in the context of the same DCR provides us with a measure of the number of initial copies of that T cell receptor present in our sample (the clone size). For this study, we additionally count the number of copies of each barcode-DCR combination (the barcode family size) to provide us with information about the amplification of the initial molecules.

The structure of the available barcode pool is inferred from the distribution of the number of times each barcode is found to have labelled a different cDNA molecule (barcode-labelling events) across all experiments in this study. The barcode-labelling events data are fitted by various zero-truncated mixed Poisson models using custom functions (found in Supplementary Information), minimised using the Optimise function of SciPy in Python. The parameters of the fitted models are used to infer the structure of the pool of available barcodes.

### PCR simulator

Simulation of labelling, amplification and sequencing of samples of molecules is performed with functions written in Python and available at github.com/uclinfectionimmunity/PCRsim. Briefly, at each cycle a molecule has a chance to successfully replicate. The probability of successful replication is determined by the PCR model chosen. If replication is successful, nucleotide error is incorporated at a given rate by choosing at random whether a given position in the sequence contains error and if so which nucleotide is incorporated incorrectly. Molecules to be sequenced are selected at random from the amplified pool and sequencing error is incorporated into these molecules similarly.

## Results

### Quantitative analysis of transcript frequency is influenced by heterogeneous amplification efficiency that is independent of target or primer sequence

**Figure 1.**
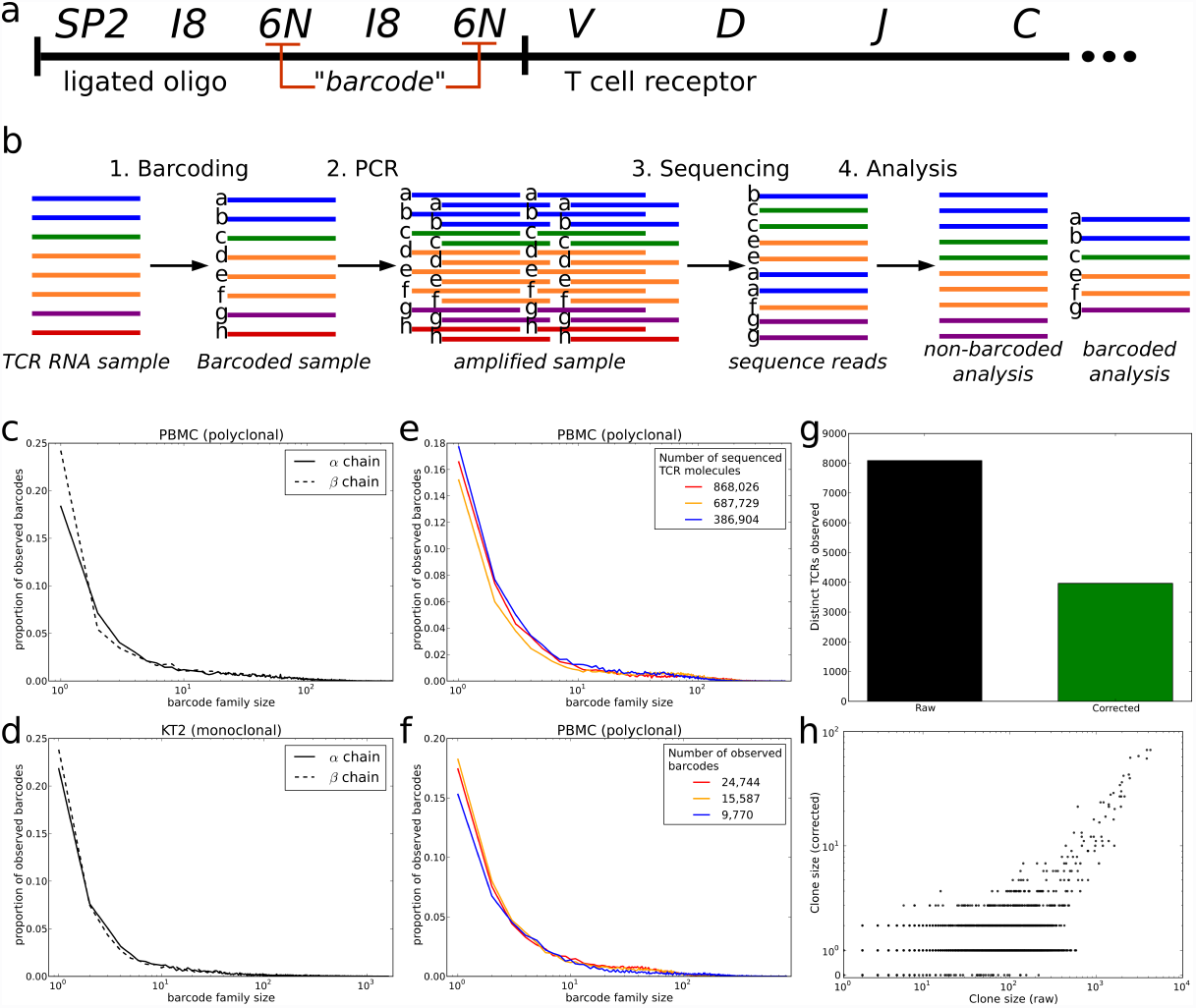
a. Schematic of the target TCR molecule with ligated oligonucleotide for molecular identification (barcoding) and sample indexing. In the T cell receptor portion of the molecule, V, D, J and C refer to the Variable, Diversity (β only), Joining and Constant regions of the TCR α or β chains respectively (not to scale). In the oligonucleotide, SP2 is the Illumina sequencing primer, I8 represents one of a set of known 8 base pair index sequences and 6N represents 6 random nucleotides. The two instances of 6N make up the 12 base pair barcode for each molecule.
b. Schematic of experimental and computational protocol used to sequence and analyse TCRs from isolated RNA. Barcodes (represented here by lower case letters) are ligated onto each TCR molecule together with a known sequence (SP2). PCR is then performed to amplify the sample. The amplified pool of molecules is diluted and introduced to the sequencer, where a sample of molecules will adhere to the flow cell and be sequenced. Repertoire analysis is performed on the sequencing data, with the barcodes allowing correction of biased PCR amplification as well as correction of sequencing errors.
c. The distribution of observed barcode family size (the number of sequence reads occurring in the sequencer output that originate from the same initial target molecule in the sample) in TCR sequence data from healthy volunteer PBMC. Solid line: TCR alpha chain data. Dashed line: TCR beta chain data.
d. The observed barcode family size distribution observed in TCR sequence data from a sample of RNA isolated from a T cell clone (KT2, responding to tetanus toxoid (30)). Solid line: TCR alpha chain data. Dashed line: TCR beta chain data.
e. The barcode family size distributions observed in TCR sequence data from healthy volunteer PB samples that are sequenced at different depths (different numbers of total sequence reads obtained).
f. The barcode family size distributions observed in TCR sequence data from healthy volunteer PB samples that contain different numbers of distinct barcodes in the sequenced sample (different number of initial molecules taken through the protocol).
g. The number of distinct TCRα clones observed in a representative sequencing run of healthy volunteer PBMC when barcodes are not considered (raw data) and when barcodes are used to error-correct and cluster the sequencer output (corrected data). Data from additional healthy volunteers shown in Supplementary Figure 2.
h. The correlation between observed clone size (the number of sequencing reads of the same TCR) in the raw data and the error- and PCR-duplication corrected data (“corrected”). Data from a representative sequencing run of healthy volunteer PBMC.

We reverse transcribed a sample of T cell receptor (TCR) RNA from PBMC, peripheral blood or a T cell clone (KT2), and then ligated a primer that contained a unique 12 base pair barcode followed by a sequence corresponding to the Illumina SP2 sequencing primer (Fig. 1a). The individually tagged mixture of different α and β chains were amplified using constant region 3’ primers and a 5’ primer homologous to the Illumina SP2 sequence on the ligated oligonucleotide. The resulting amplified PCR reaction was diluted and sequenced using the standard Illumina protocol (Fig. 1b). The number of times each barcode was present in the sequence data was then counted. We refer to all sequences that have an identical barcode as a barcode family, and refer to the number of molecules present with this barcode as a barcode family size. Although each cDNA molecule was ligated to a different barcode, and the starting frequency of each barcode should then be uniform and independent of the frequency of the TCR sequence with which it was associated, the observed distribution of barcode family sizes in a PBMC sample was very heterogeneous (Fig. 1c). Thus, while the majority of barcode families were of size one, some barcodes occurred over 100 times. A similar pattern was observed for α and β TCR sequences, indicating that the heterogeneity was not some special feature of the sequence being amplified.

One obvious explanation was that, although the primers and the primer targets were the same for all amplified molecules, the intervening sequences were heterogeneous since they represented many different TCR sequences. Thus, heterogeneous amplification could reflect differences in target replication by polymerase. In order to rule out this trivial explanation, we labelled and amplified a single TCR sequence (α and β chain) from a human T cell clone, KT2. As predicted, the vast majority of sequences from these samples were identical, and the rest probably represented PCR or sequencing error (Supplementary Fig 1). To our surprise the distribution of barcode frequencies was still just as heterogeneous (Fig. 1d). Thus even under conditions where target, primer and reaction conditions were identical for all amplified molecules, we observed a very significant heterogeneity in apparent amplification efficiencies. We repeated this experiment using different PBMC samples, sequenced at different depths (Fig. 1e) and with different numbers of initial target molecules (therefore different numbers of observed barcodes) carried through the protocol (Fig. 1f) and obtained similar results.

**Figure 2.**
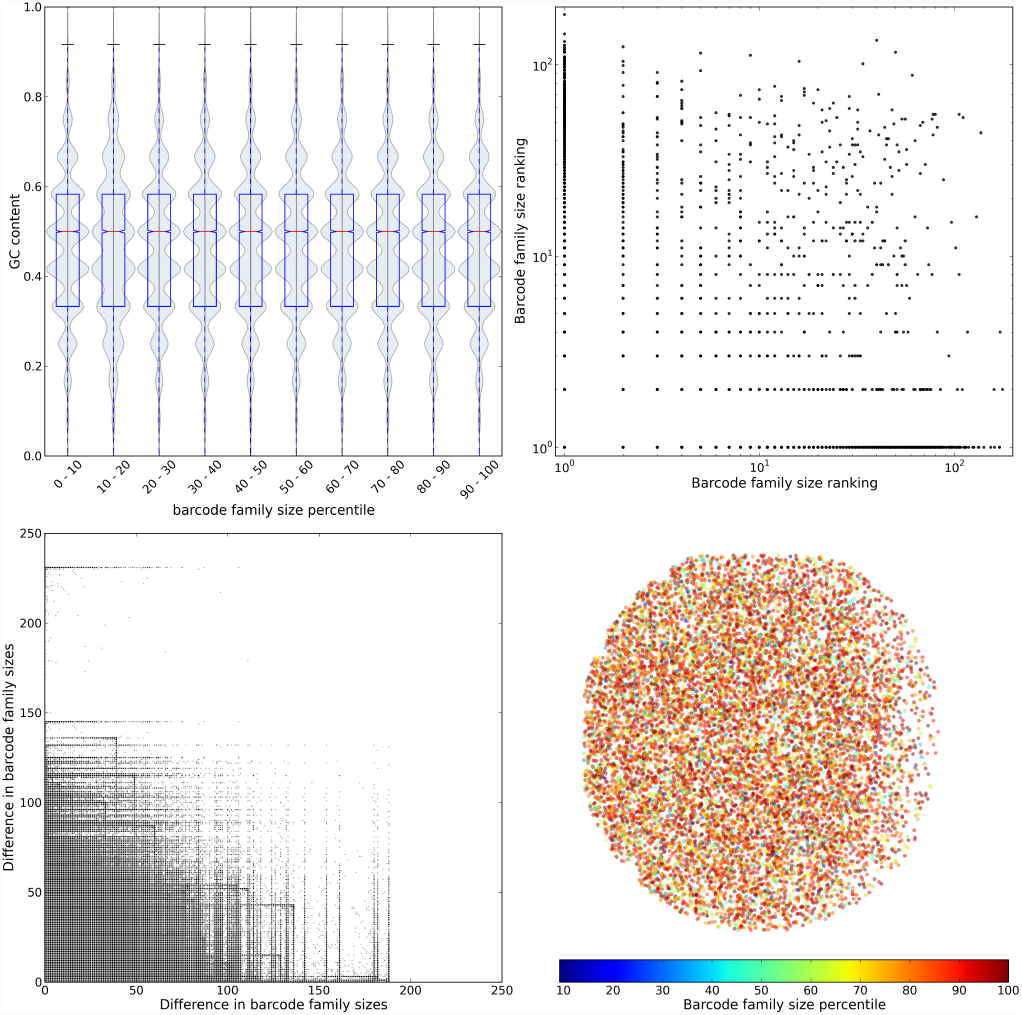
a. Distribution of GC content of the 12-nucleotide random barcodes by barcode family size percentile. Data from a sequence run of healthy volunteer PBMC TCRs.
b. The correlation between the barcode family size ranking in any pair of runs for those barcodes that occur in more than one of the eight monoclonal KT2 TCR sequencing runs in this study (R-squared < 0.0003). Ranking is ascending, and barcodes that have the same family size in a run are given the same ranking. There is no gap introduced in rankings when more than one barcode occupies a particular ranking, as such for small barcode family sizes ranking is equivalent to barcode family size.
c. For those pairs of barcodes that appear together in any pair of the eight KT2 sequencing runs in this study, the relationship between the difference in barcode family sizes in one run and in the other. R-squared < 0.0004.
d. Position of TCR molecules on the flowcell, coloured by barcode family size percentile. Representative example of a single frame from one flow cell from a sequencing run in this study.

It is important to note that this amplification heterogeneity, although it is unrelated to the biological heterogeneity in TCR sequences (the repertoire), nevertheless materially affects quantitative features of the observed repertoire. Sequencing TCRs with our barcoding technique allows correction of PCR amplification error, by counting the number of different barcodes appearing with the same TCR, as well as correction of sequencing error, by comparing sequence reads from the same barcodes. Correcting for PCR- and sequencing-error in the sequencing data significantly reduces the apparent diversity of the observed TCR repertoire (Fig. 1g, Supplementary Fig 2). In addition, although large TCR clones are observed many times in both raw and corrected data, there is a lot of noise introduced by heterogeneous amplification in the smaller clones if barcoding is not incorporated into the analysis (Fig. 1h, Supplementary Fig 2).

### Barcode family size is not dependent on barcode sequence, barcode clash or nonCuniform barcode primer frequencies

The heterogeneous amplification observed could hypothetically be caused by the barcode itself since the polymerase must amplify the barcode in each cycle. To investigate this, we first considered whether barcodes that appear more amplified have a tendency to contain more or fewer G or C nucleotides (Fig. 2a). However, there was no obvious relationship between the frequency of particular barcodes and their GC content. Furthermore, the frequency of the same barcode in any two different sequence runs was uncorrelated (Fig. 2b). A high barcode family size did not therefore appear to be the result of a particular barcode sequence or sequence motif. Additionally, to account for the fact that the amplification effect might be to do with relative, rather than absolute, barcode ‘fitness’, we considered all pairs of barcodes that both appear in any pair of experiments. If the amplification was determined by the barcode we would expect, for example, that if barcode A is larger than barcode B in experiment 1 then it would also be larger in experiment 2. We found no correlation between the frequencies of any two barcodes that appear together in a pair of experiments (Fig. 2c), implying that the barcode sequence itself does not determine the efficiency with which each molecule is amplified. We also examined whether the observed barcode family size might be an artefact introduced during the sequencing reactions, perhaps by heterogeneity in bridge PCR on the flow cell. If this were the case we would expect that molecules from large barcode families are located in close proximity on the flow cell. However there was no observable relationship between barcode family size and location of molecules on the flow cell (one representative frame shown in Fig. 2d).

**Figure 3.**
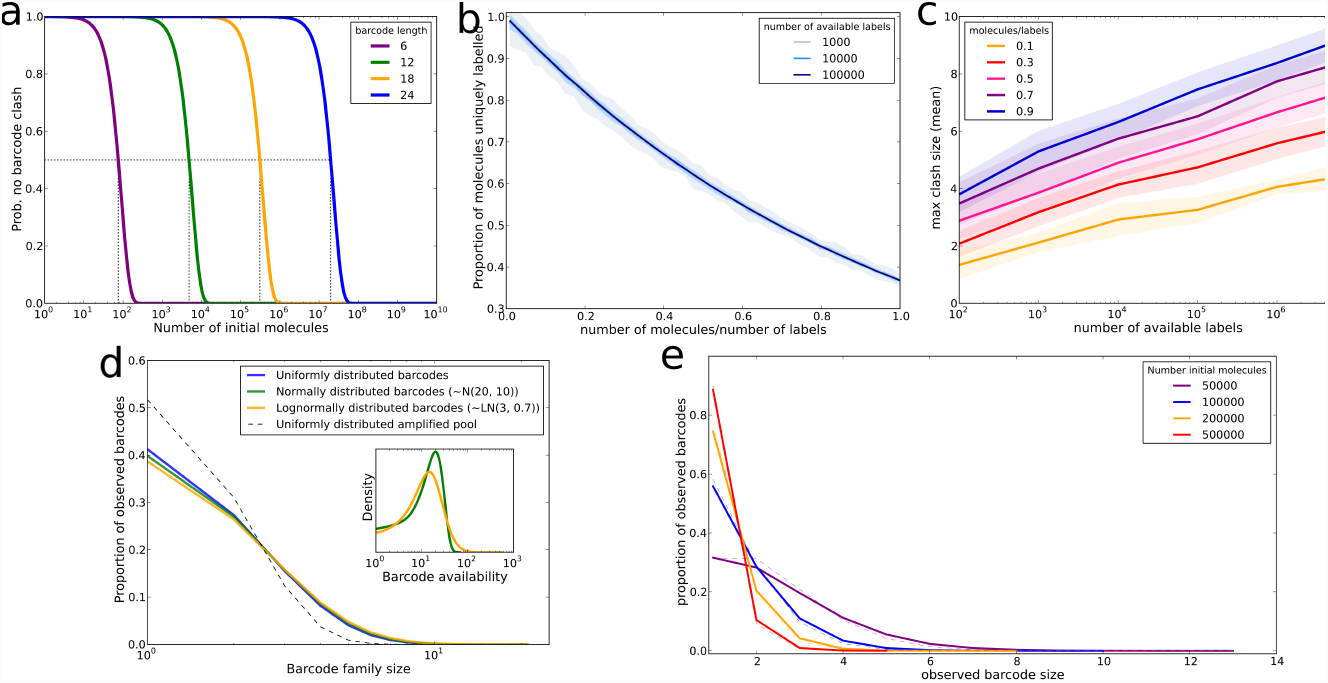
a. The probability that there is no two molecules receive the same label (“barcode clash”) when initial molecules are labelled with a pool of random nucleotide barcodes of the indicated length. The dotted lines indicate the number of molecules that can be labelled with a 50% chance of no barcode clash occurring.
b. The proportion of initial molecules that receive a unique barcode when barcoding is simulated with the indicated number of available barcodes, uniformly distributed. The number of molecules to be barcoded is expressed as a proportion of the number of available barcodes. Data shown is the mean and standard deviation of 50 repeated simulations.
c. The maximum number of initial molecules that receive the same barcode when barcoding is simulated with the indicated number of available barcodes that are uniformly distributed. The number of molecules being barcoded is indicated by colour, expressed as a proportion of the number of available barcodes. Data shown is the mean and standard deviation of 50 repeated simulations.
d. The observed family size distribution after simulation of barcoding 250,000 initial molecules from uniformly or non-uniformly distributed pools of 500,000 available barcodes, 10 cycles of PCR at efficiency 0.5 and sequencing 300,000 molecules from the amplified pool. The distribution of available barcodes for the non-uniform simulations is shown in the inset (green: normal distribution (restricted to values > 0), orange: lognormal distribution). Data shown are the mean and standard deviation of 10 repeated simulations. The grey dotted line shows the barcode family size distribution that would be expected if the molecules to be sequenced were drawn from a uniformly distributed amplified pool, in which all molecules had been uniquely barcoded and amplified equally.
e. The observed barcode family size distribution when the indicated numbers of initial molecules are barcoded from a pool of 412 potential barcodes with barcode availability distributed as predicted from empirical labelling events observed (details in supplementary information, empirical distribution shown in Supplementary Figure 3). PCR cycles (25 cycles, 0.75 efficiency) are simulated on the labelled molecules, and samples of size 100,000 are selected from the amplified pool. The solid line represents the mean of 10 repeated simulations. The dashed line shows the expected distribution had the sample been drawn from a uniformly distributed amplified pool, in which every initial molecule had been barcoded uniquely and amplified by the same amount.

The barcodes should theoretically contain randomly chosen nucleotides at each of the 12 positions, giving a total of 4^12^ ≈ 1.7 × 10^7^ possible barcodes, each appearing an equal number of times. In practice, the methods of oligonucleotide synthesis likely result in slightly different incorporation efficiencies of different nucleotides at each position (27). In addition, the number of target molecules barcoded in our T cell samples is often within an order of magnitude of the number of available barcodes, resulting in a significant probability that the same barcode is used more than once (‘barcode clash’) (Fig. 3a). In order to assess the impact that this barcode clash might have on the observed barcode family sizes, we first simulated barcoding molecules from a large, uniformly distributed pool of available barcodes and measured the proportion of molecules that were uniquely barcoded (Fig. 3b). This value depends on the ratio of the number of available barcodes (size of the barcode pool) to the number of molecules to be barcoded. In these simulations we also measure the maximum observed barcode clash size (Fig. 3c), which in contrast also depends on the absolute number of available barcodes and molecules to be barcoded. These simulations show that in our protocol (barcoding in the order of 10^6^ molecules with 10^7^ available barcodes) around 90% of molecules get a unique barcode and the maximum clash size is predicted to be below 4. Thus barcode clash is unable to account for the range in barcode family sizes we observe in our data.

It is likely that the pool of barcodes we have available for labelling is not exactly uniformly distributed, which could lead to increased barcode clash. We simulated the barcoding, amplification and sequencing protocol using normally or lognormally distributed barcode frequency distributions, but this had little effect on the observed barcode family size distributions when compared to uniquely barcoding every molecule or to the expected distribution if every initial molecule was represented equally in the post-PCR amplified pool (Fig. 3d). We also derived the empirical distribution of barcodes in our initial oligonucleotide pool (see Supplementary Information and Supplementary Figure 3) and simulations using this distribution do not show a barcode family size distribution deviating far from the sampling distribution expected from a uniformly distributed amplified pool (Fig. 3e). The output of the barcoding, amplification and sequencing pipeline is therefore robust to the likely occurrence of barcode clash and non-uniform barcode frequencies.

### Inherited differences in PCR efficiency are necessary to explain the observed diversity in barcode family size

**Figure 4.**
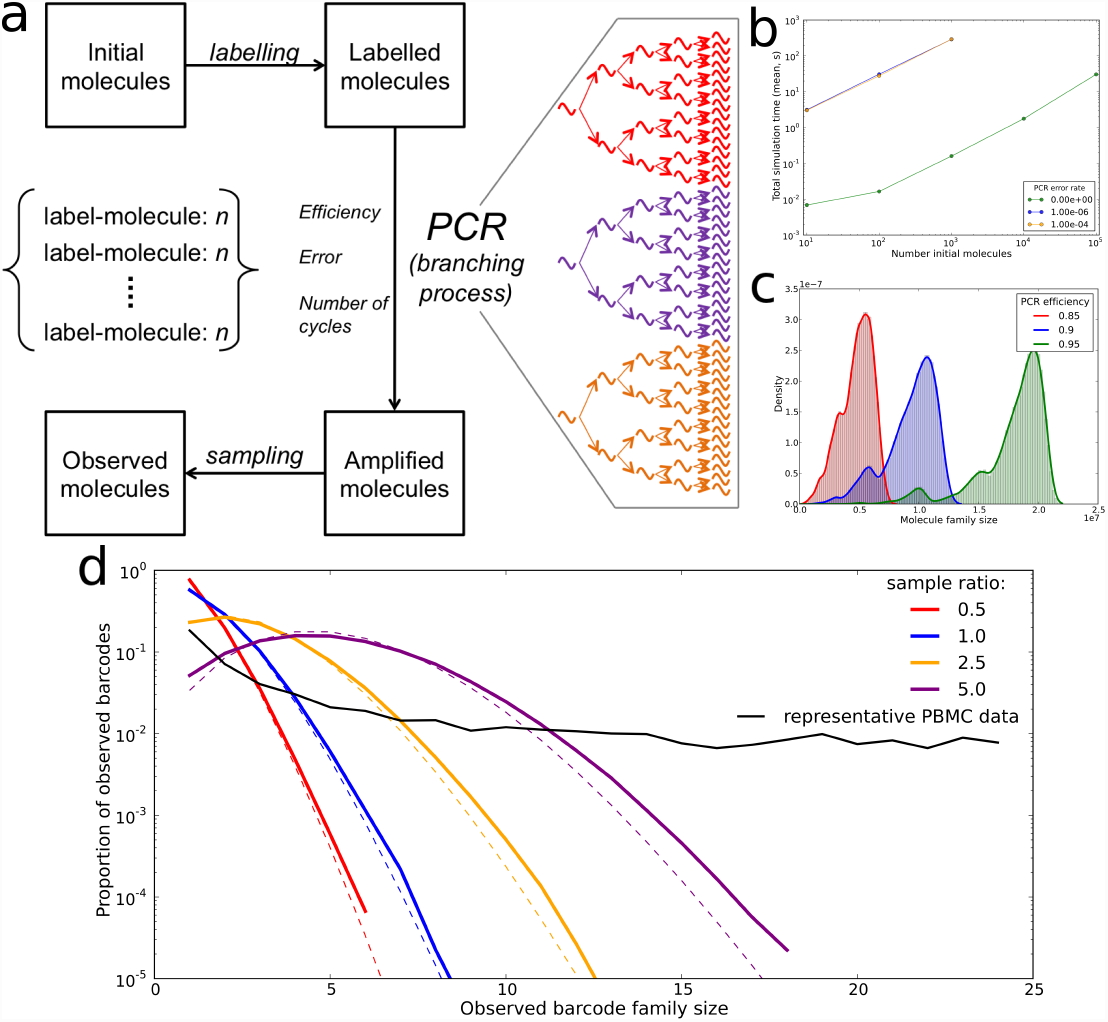
a. Schematic of the PCR simulator software used in this study. The software includes adding barcodes to molecules (‘labelling’), PCR amplification with a specified number of cycles, efficiency model and error rate, and sampling and sequencing from the amplified pool.
b. Time taken to perform a full simulation, which includes initialisation, labelling initial molecules, PCR cycles (using a standard branching process model), sampling from the amplified pool and sequencing. Simulations are performed with the indicated PCR error rate (per base per cycle) and the given number of initial template molecules. Simulations consist of 15 cycles of PCR with efficiency 0.8, a sample size equal to the number of initial molecules being chosen from the amplified pool and sequencing with error rate 10-4. Data shown is the mean of 5 repeated simulations at each set of conditions, as measured on a 2.8 GHz Intel Core i7 MacBook Pro.
c. The distribution of the number of copies of each of 100,000 initial target molecules after 25 cycles of PCR at efficiencies of 0.85 (red), 0.9 (blue) or 0.95 (green).
d. The distribution of observed barcode family sizes (coloured lines) after simulating PCR cycles (25 cycles at 0.9 efficiency) on 100,000 initial molecules and then sampling from the amplified pool to select those molecules that are observed in the sequencer output. The number of molecules sequenced is expressed as a proportion (the ‘sample ratio’) of the number of initial molecules (100,000). The solid coloured lines are the mean of 5 repeated simulations, and the dashed coloured lines are the expected distribution (a zero truncated Poisson with parameter equal to the sample ratio) if the sample was drawn from a uniformly distributed pool (which would occur if every initial molecule was uniquely barcoded and amplified identically). The black solid line is a representative example of the barcode family size distribution observed in TCR sequencing data from healthy volunteer PBMC.

The experimental pipeline involves amplification followed by subsampling for sequencing, which can introduce Poisson non-uniformity even when the amplified pool of barcoded molecules is uniform. Furthermore, PCR efficiencies of less than 100% can introduce non-uniformity resulting from the inherent stochasticity of the PCR process (28). In order to examine how variable efficiency and sampling could affect observed barcode family size distributions we developed a PCR simulator in which molecules are barcoded, amplified and then sampled in silico. The simulator is outlined schematically in Figure 4a. In its most basic implementation (modelling PCR as a straightforward branching process with no error) the simulator can perform a full simulation (labelling initial molecules, performing 15 PCR cycles with efficiency 0.8, sampling and sequencing including sequencing error) on 105 initial molecules in approximately 12 seconds (Fig. 4b). Introducing PCR error substantially increases the simulation time, although altering the error rate further does not alter simulation time. Parallelisation and cluster research computing platforms make PCR simulation including error of large numbers of initial molecules feasible.

The simulated barcode distributions (the number of molecules present after amplification that are derived from each initial molecule) at different efficiencies are shown in Figure 4c. The introduction of less than 100% efficiency introduces some barcode family size heterogeneity as described previously (28). This variation arises because, in every replication cycle, any individual molecule may or may not replicate with a probability determined by the overall efficiency. The substantial shoulder observed in the distributions correspond to molecules which fail to be replicated in the first cycle of PCR and hence are present at half the average number of copies. However, the heterogeneity caused by low efficiencies is averaged out over many molecules and the majority of barcode family sizes are within a factor of two of each other at the end of the PCR reaction.

When a sample of molecules is drawn at random from the amplified pool (to simulate the process by which molecules from the amplified sample are diluted and introduced to the flow cell to anneal to complementary capture oligonucleotides), the observed barcode family size is further diversified depending on the ratio of number of sequenced molecules to number of initial molecules (Fig. 4d). These observed barcode family size distributions follow a Poisson distribution (as an approximation to a binomial distribution), scaled to account for the fact that we cannot count those barcodes with an observed family size of zero (a zero-truncated Poisson). The Poisson distribution is the expected distribution when sampling from an amplified pool in which each barcode is present the same number of times. If the PCR process in our experiments behaved as a straightforward branching process we would expect our experimental observed barcode family size distributions to also follow a zero-truncated Poisson distribution, with the Poisson parameter providing information about how many initial molecules there were in our sample. However, it can be seen that our data does not belong to the same family as the simulated distributions (Fig. 4d), suggesting that these samples were not drawn from a uniformly distributed post-PCR pool and that neither low PCR efficiency or the sampling process can account for the broad distribution observed.

We therefore tried to formulate variations of the branching process model of PCR that could explain the broad barcode family size distribution observed.

**Figure 5.**
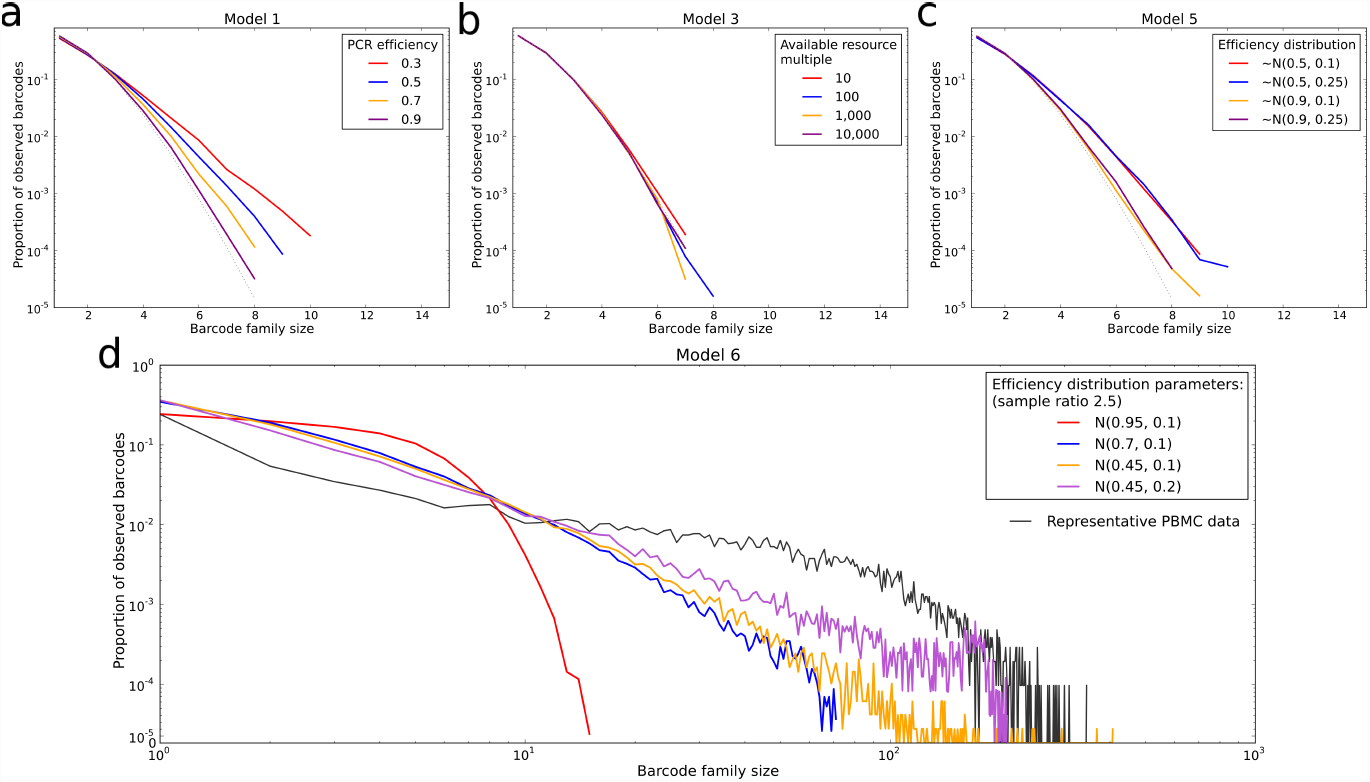
(a) – (c) Observed barcode family size distributions observed under different models of PCR duplication. Simulations performed with 10,000 initial molecules, 25 cycles of PCR (with no error) and sequencing of 10,000 molecules selected from the amplified pool. Simulations were repeated 10 times and the mean and standard deviation are shown. The dotted lines represent the expected distribution if every initial molecule is barcoded uniquely and represented equally in the amplified pool. Models described in the text but not displayed here can be found in Supplementary Figure 4.

a. Model 1: Standard branching process of PCR, with the indicated efficiencies. The efficiency is the probability that a given molecule will duplicate in a given cycle.
b. Model 3: Model of PCR where the duplication efficiency depends on competition between target molecules for a constant level of resource, given as a multiple of the number of initial molecules.
c. Model 5: Variable efficiency model of PCR, where the probability of a given molecule replicating in a given cycles is selected from a normal distribution (restricted to [0, 1]) with the indicated parameters (mean and standard deviation).
d. Model 6: Inherited efficiency model of PCR, where the probability of replication in a given cycle is identical for all molecules derived from the same initial molecule. The efficiencies for the initial molecules are selected from a normal distribution with the indicated parameters (mean and standard deviation). 25 PCR cycles are simulated on 10,000 initial molecules, and then a sample is drawn from the amplified pool at a multiple of 2.5 times the number of initial molecules. The observed barcode family size distribution shown is the mean of 10 repeated simulations. In black is a representative barcode family size distribution from a healthy volunteer PBMC sample.

The starting point is a standard branching process model of PCR (‘model 1’) where the efficiency of the PCR (between 0 and 1) refers to the probability that a molecule will replicate successfully in a cycle. Using this model, we simulate PCR and sampling, and show that the resulting barcode family size distributions do not diverge significantly from the expected Poisson distribution regardless of the efficiency used (Figure 5a). Next, a target degradation model (‘model 2’) was used. Model 2 is set up as for model 1, except that when a molecule fails to duplicate in a cycle there is a chance that it instead degrades and is no longer available to be amplified in later cycles of the PCR. Again, simulation of this model does not reproduce the large deviation from the expected Poisson seen in our data (Supplementary Figure 4a).

Next, we introduce competition for resource, which affects the success rate of duplication of molecules. This abstract ‘resource’ covers, for example, the availability of dNTPs and primer in the PCR mixture, and the ability of the enzyme to process the molecules inside the time frame given in the PCR protocol. The first resource competition model (‘Model 3’) is one in which there is a fixed, constant amount of resource available and the probability that a molecule successfully replicates in a cycle is given by the number of molecules present at the start of the cycle divided by the amount of resource. As such, the efficiency of the reaction decreases through the cycles once the number of molecules present exceeds the capacity of the available resource to process all those molecules in one cycle. Figure 5b shows that this model cannot reproduce the spread of barcode family sizes we observe in the data. An alternative resource competition model (‘Model 4’) involves degradation of resource as it is used, at a given degradation rate. This model is also unable to account for our observed barcode family size distributions (Supplementary Figure 4b).

Instead of a constant efficiency across all molecules and all cycles, we imagine that in a given cycle some molecules are able to replicate more efficiently than others. For instance this variation may depend on the position of the molecule within the sample (which may affect e.g. proximity to primer) or the conformation of the molecule (which may affect ability of the primer to bind). We introduce a variable efficiency model (‘Model 5’), where the probability that a given molecule will replicate in a given cycle is chosen from a defined distribution. Model 5 is implemented using a normal distribution with a variety of parameters (Figure 5c). Although a low mean efficiency and a large standard deviation produces the most divergence from the expected barcode family size distribution, none of the parameters investigated was able to reproduce the observed spread of family sizes.

We adapted Model 5 to include the constraint that once an efficiency is chosen for a molecule in cycle 1 this same efficiency is inherited by all molecules produced from this initial molecule (‘Model 6’). Simulation of PCR and sampling using model 6 was performed, and showed that inherited efficiencies could produce a substantial amount of spread in the barcode family size distribution when the efficiency distribution has a low mean and a relatively large standard deviation (Figure 5c). The observed barcode family size distribution from Model 6 can be seen to be broadly comparable to that seen in our experimental data.

## Discussion

PCR is a fundamental and ubiquitous tool of molecular biology laboratories. The combination of PCR and HTS, in particular, has driven an explosion in DNA sequence acquisition. In many of these applications, for example RNA-seq and lymphocyte antigen receptor repertoire studies, the quantification of transcripts is critical, since the output is based on counts of specific sequences. The avoidance of PCR bias is therefore critical and much effort has been expended on trying to control and mitigate bias. In this study, we examine the consistency of PCR amplification, using molecular barcodes to follow amplification of single molecules. We find that the distribution of the number of copies of an initial molecule that are observed in sequencer output varies over a wide range, in a manner which is independent of primers, target sequence, bulk PCR conditions or barcode sequences.

The differential binding properties of different primers, and secondary structure within target sequences are well-established causes of PCR biases. Multiplex PCRs, for example, frequently show different efficiencies for different primer/target combinations. This bias is a known confounder of T cell repertoire studies, for example. As a result, we and others (16) have developed techniques that use various types of 5’ RACE, and thus can amplify with amplicon-independent primers. However, the variation in target sequence to be amplified is obviously a variable that cannot be avoided. In this study we therefore consider the extent to which amplification bias can be attributed to sequence variability. We compare the amplification of heterogeneous mixtures of alpha or beta T cell receptor chains (typically containing >10^4 different sequences) with amplification of a monoclonal T cell receptor from a T cell clone (this clone in fact expresses more than one TCR chain, a common feature of T cells (29)). Unexpectedly, PCR amplification efficiency (the number of observed molecules derived from a single ancestor) varies broadly, both for the polyclonal and monoclonal populations. Indeed the extent of variability is very similar, suggesting that the actual sequence of the TCR variable region is not the major cause of different amplification rates. Our results do not, of course, imply that all sequences will be amplified equally. Indeed the length of the target and the GC content are well known to influence PCR efficiency (6). Rather our results suggest that even when amplifying relatively small amplicons (<1KB) whose sequences are all rather comparable, substantial variation remains.

If the variability is not due to primers or target sequence what might be the cause? Our data suggest that the sequence of the ligated barcodes is not the cause of the observed differential amplification, since barcode family size is not correlated between experiments. Indeed it seemed a priori unlikely that if the variation cannot be attributed to differences between V region sequences, it would result from 12 base pair barcodes. Additionally, analysis of the structure of the pool of the random barcodes that are used to label the initial molecules suggests that while there is potential for barcode ‘clashes’ (where the same barcode is chosen to label more than one initial molecule), these are not large enough or prevalent enough to be the reason for the large barcode family sizes observed. We do, however, present some theoretical and simulation results that can help to guide the size of barcode pool size in different scenarios. These results suggest that a barcode of 12 base pairs (providing in the order of 10^7^ different sequences) is sufficient to label pools of DNA targets in the order of 10^6^ molecules.

The bulk conditions in all the PCR reactions obviously cannot account for the intra-experimental variation. However, as discussed previously, PCR is by its nature a stochastic process since at each cycle a molecule will be either replicated or not replicated with some probability p, which will be less than 1 for all reactions in which replication efficiency is not 100%. For example, the PCR efficiencies in our model system (which we have measured using qPCR on plasmid dilutions) are typically in the order of 1.8-1.9. Furthermore, it is possible that there is local heterogeneity in the PCR vessel itself: for example temperature gradients, or heterogeneity introduced by phase shifts at the plastic/liquid or liquid/gas surfaces. We therefore examined the implications of these different models in detail using a branching process PCR simulator.

Simulation demonstrated clearly that lower efficiencies, a range of efficiencies, competition and resource limitation can all introduce some variation in the predicted output of the PCR for different molecules. As might be predicted, the extent of variation increases with cycle number, and with low and more variable efficiencies. The goal of minimising the number of cycles, and maximising efficiency does therefore lower overall expected variance of product molecular counts. However, the extent of the variance is limited and does not explain our observed results. The only model we considered that was able to produce substantial variance in output comparable to that observed is an inherited efficiency model, where all molecules produced from an initial molecule retain the same efficiency though all cycles. This result, too, is related to well known evolutionary theory where significant divergence can only occur when selection operates on the inherited properties of the individual. The nature of these properties, in the case of PCR, remains an unanswered puzzle. We have ruled out that they could arise from the sequence itself, the most obvious explanation. One can speculate that perhaps some DNA molecules are trapped at the plastic surface of the PCR tube, or at the water/air interface by local surface tension effects and that these molecules are replicated differently from the bulk molecules in solution.

In conclusion, we consider the implications of our findings for the community routinely using PCR for quantitative analysis of RNA or DNA populations. The major lesson is that molecular barcoding provides an essential tool that can mitigate for the effects of PCR heterogeneity. This is especially important for studies whose primary output is the comparative quantification of many diverse nucleotide fragments within a mixture, such as repertoire analysis. In situations where single molecule barcoding is difficult, or not practical, every effort needs to be taken to maximise the efficiency of the PCR reactions and minimise the number of cycles. In the longer term, single molecule amplification-free DNA sequencers, which are currently in development, may remove the requirement for a PCR amplification step altogether. In the meantime, it continues to be important to appreciate the inherent stochasticity of the PCR process, and its possible effects on quantitative aspects of molecular biology.

## Author Contributions

KB and JH performed analysis of the data. KB and BC wrote the manuscript. TO, JH and BC developed the experimental protocol. All authors reviewed the manuscript.

## Competing financial interests

The authors declare no competing financial interests.

## Funding

This work was supported by Engineering and Physical Sciences Research Council and Medical Research Council. Theres Oakes is supported by a grant from Unilever plc.

